# Derivation, Validation, and Prediction of Loading-Induced Mineral Apposition Rates at Endocortical and Periosteal Bone Surfaces Based on Fluid Velocity and Pore Pressure

**DOI:** 10.1101/2023.09.08.556956

**Authors:** Sanjay Singh, Satwinder Jit Singh, Jitendra Prasad

## Abstract

Bones adapt to mechanical forces driven by osteocytes in the lacunae canalicular network (LCN). Osteocytes sense signals such as strain, interstitial fluid flow, and pore pressure. Physiological tissue strain is insufficient to induce bone formation, and fluid flow-based models struggle to predict bone formation at both the periosteal and endocortical surfaces, simultaneously. This prompted the exploration of pore pressure’s role.

The study introduces dissipation energy density as a more significant stimulus, combining various LCN parameters, including the waveforms of both fluid velocity and pore pressure and the number of loading cycles. A pivotal achievement is the mathematical derivation of the Mineral Apposition Rate (MAR), linked proportionally to the square root of the dissipation energy density minus its reference value. This hypothesis is subjected to testing/ validation for both endocortical and periosteal surfaces for an in-vivo study on mouse tibia available in the literature.

Computational implementation of this mathematical model adopts a poroelastic finite element analysis approach, treating bone as a porous fluid-filled entity. Crucially, assumptions underpinning the model, such as the impermeability of the periosteum to fluid and the maintenance of a reference zero pressure at the endosteal surface, are corroborated by relevant experimental studies.

As a bottom line, the resulting model is the first of its kind, as it has been able to predict MAR at both endocortical and periosteal surfaces, significantly advancing our understanding of cortical bone adaptation to exogenous mechanical loading.

## 1 Introduction

The investigation into bone adaptation to mechanical forces dates back to Von Meyer and Karl Culmann, who proposed the alignment of trabeculae with stress trajectories [1]. Three papers in the late 1900s paved the way for load-induced fluid flow in LCN as the primary stimulus for bone adaptation [2-4]. Weinbaum et al. [2] hypothesized that fluid shear on osteocyte processes in LCN initiates a cellular response. Cowin et al. [3] suggested LCN as the site of the strain-generated potential. Klein-Nulend et al. [4], in their in vitro study, showed osteocyte sensitivity to fluid flow shear stresses. Based on these theories, fluid flow is believed to predict new bone formation, and accordingly, several fluid flow-based computational models emerged. For instance, Kumar et al. [5] devised a viscous dissipation energy-based mathematical model to predict new bone formation. However, the work was based on a solid rectangular beam, not a real bone. There was no endocortical surface, and no validation done with respect to experimental data. Dissipation energy density used in that, however, proved very useful and intuitive to incorporate almost all types of mechanical stimuli. Periera et al. [6] introduced the fluid velocity-based mathematical model to anticipate changes in cortical thickness which does not provide quantitative details of the new bone formation individually at the periosteal and endocortical surfaces. Carriero et al. [7] tried to correlate the peak fluid velocity to the new bone formation; however, the peak fluid velocity and the new bone formation do not coincide at the endocortical surface. On the other hand, periosteal surface bone formation was greatly overestimated, which (according to authors) was a result of using the high permeability value of the periosteum in accordance with the ex-situ measurement of permeability by Evans et al. [8], who had also found the periosteum permeability to be stress-dependent and direction dependent. Therefore, it is also difficult to implement this highly nonlinear behavior of the periosteal permeability in a bone adaptation model. Moreover, comparing the results of the in-situ experiment of Price et al. [9] to that of the corresponding theoretical/computational model of Zhou et al. [10], the complete impermeability of the periosteal surface seems to be plausible.

Tol et al. [11] also developed a fluid velocity-based theoretical model. Here also, new bone formation is not found to vary monotonically with the fluid velocity at the periosteal and endosteal surfaces. To our knowledge, there is no example in the literature where both endocortical and periosteal bone surfaces can be predicted simultaneously by fluid flow alone. It is, therefore, essential to understand the mechanical environment of osteocytes inside the LCN and re-analyze it.

Osteocytes residing in LCN are enveloped by fluid and anchored to mineralized tissue through tethering fibers. Additionally, these processes are connected to the projection protruding from the canalicular wall through *α*_*v*_ *β*_3_ integrin molecules, as depicted in Fig. 1(a) [12]. Tissue strain in the whole bone under the physiological loading is typically less than 0.2% [13, 14]. However, this strain must be an order higher for any intracellular response to occur [15]. When the mechanical load is applied to the bone, it induces a pressure gradient, leading to canalicular fluid flow. This fluid flow, in turn, exerts drag forces on these tethering fibers. This drag stretches the osteocyte’s process membrane outward, resulting in radial strain, leading to the deformation of the canalicular process, as illustrated in Fig. 1(b) [16]. Intuitively, pore pressure is also expected to radially stretch/compress the osteocyte’s process, depending on the tensile/compressive environment. Thus, osteocytes will sense the combined effect of fluid flow and pore pressure. The multitude of in vitro studies confirm that fluid flow stimulation prompts osteocytes to release biomolecules (Prostaglandin E_2_, Ca^2+^, etc.), culminating in osteogenesis [4, 17, 18]. Some in vitro studies demonstrate that besides interstitial fluid flow, cyclic pore pressure also has the potential to induce osteogenesis [4, 17, 19, 20]. Weinbaum et al. [2] and Scheiner et al. [21] established that fluid flow on the osteocyte process and pore pressure generated under physiological loading conditions are adequate for osteocyte stimulation [21]. Given the limitation of fluid flow alone, investigating a combination of fluid flow and pore pressure became imperative.

**Fig. 1.**
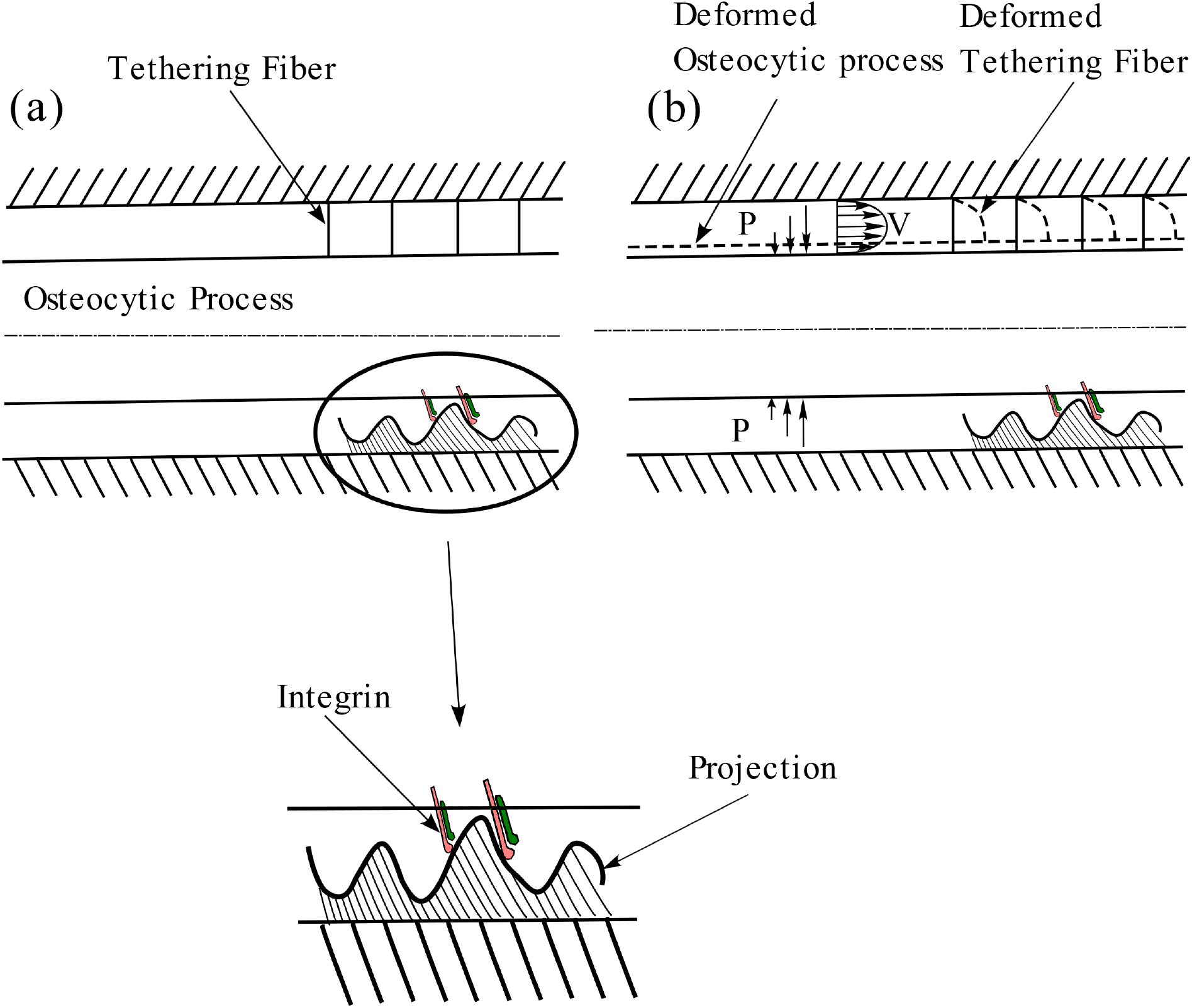
Schematic diagram of longitudinal osteocyte process and attachments, (a) before and (b) after deformation (P and V represent Pressure and Velocity of fluid).

A combination of pore pressure and fluid flow is especially intuitive and important given that the periosteal bone surface is impermeable, so fluid flow is minimal at the periosteal surface. Hence, something other than fluid flow must induce new bone formation there. On the other hand, pressure is very high at the periosteal surface due to impermeability; hence, consideration of pressure makes sense. With these factors in mind, we explore the role of interstitial fluid flow and fluid pore pressure in osteogenesis.

This study also introduces an innovative derivation of Mineral Apposition Rate (MAR) in terms of dissipation energy, which incorporates both fluid flow and pore pressure, along with the number of bouts of loading. The derivation suggests that MAR is proportional to the square root of dissipation energy density minus a reference value, forming the central hypothesis to be verified. To test this hypothesis, the dissipation energy density is computed based on the results (Fluid velocity and Pore pressure) of the Finite Element Analysis of the mouse tibia, which has been modeled as a poroelastic material. Mathematical model parameters are computed by fitting data to an experimental study on periosteal and endocortical surfaces. The model successfully predicts the MAR obtained by the experiment at both endocortical and periosteal surfaces. In addition, the model also successfully predicts new bone formation for a different condition.

The work advances the understanding of load-induced bone adaptation by establishing: (i) Pore pressure can also induce new bone formation, unlike the existing notion that fluid flow alone causes new bone formation, (ii) Pore pressure in combination with fluid velocity is a better predictor of MAR than fluid velocity alone, (iii) Dissipation energy density is a suitable stimulus as it can combine different types of stimuli such as fluid velocity and pore pressure into one scalar value. (iv) MAR is proportional to the square root of the dissipation energy minus a reference value. (v) Assumption that the Periosteal surface is impermeable is valid not only based on literature but also based on the outcomes of the developed model.

The rest of the work is organized as follows. Section 2 presents the methodology used in this work, including the derivation of MAR as a function of dissipation energy density, the finite element model used, computation of dissipation energy density due to fluid velocity and pore pressure, determination of model parameters fitting the experimental data, and statistical analysis. Section 3 presents the results. The sensitivity analysis of a model parameter has been carried out in Section 4. The results and the developed model have been discussed in Section 5 A conclusion has been made in Section 6.

## 2 Methods

### 2.1 Finite element model

We used the poroelastic finite element model to compute fluid velocity and pore pressure in bone for a given loading. The commercial ABAQUS software (Dassault Systèmes Simulia Corp) was used for this purpose.

#### 2.1.1 Bone Geometry

Analysis of the full bone would have been ideal. However, the poroelastic modeling of a full bone is computationally very costly in terms of the simulation time. In view of this, as shown in Fig. 2, we used a prismatic beam of 5 mm in length with a cross-section similar to that of a tibial mid-diaphysis located at 1.8mm proximal to the tibia-fibula junction of an 8-week-old mouse. The cross-section has been adapted from Berman et al. [22].

**Fig. 2.**
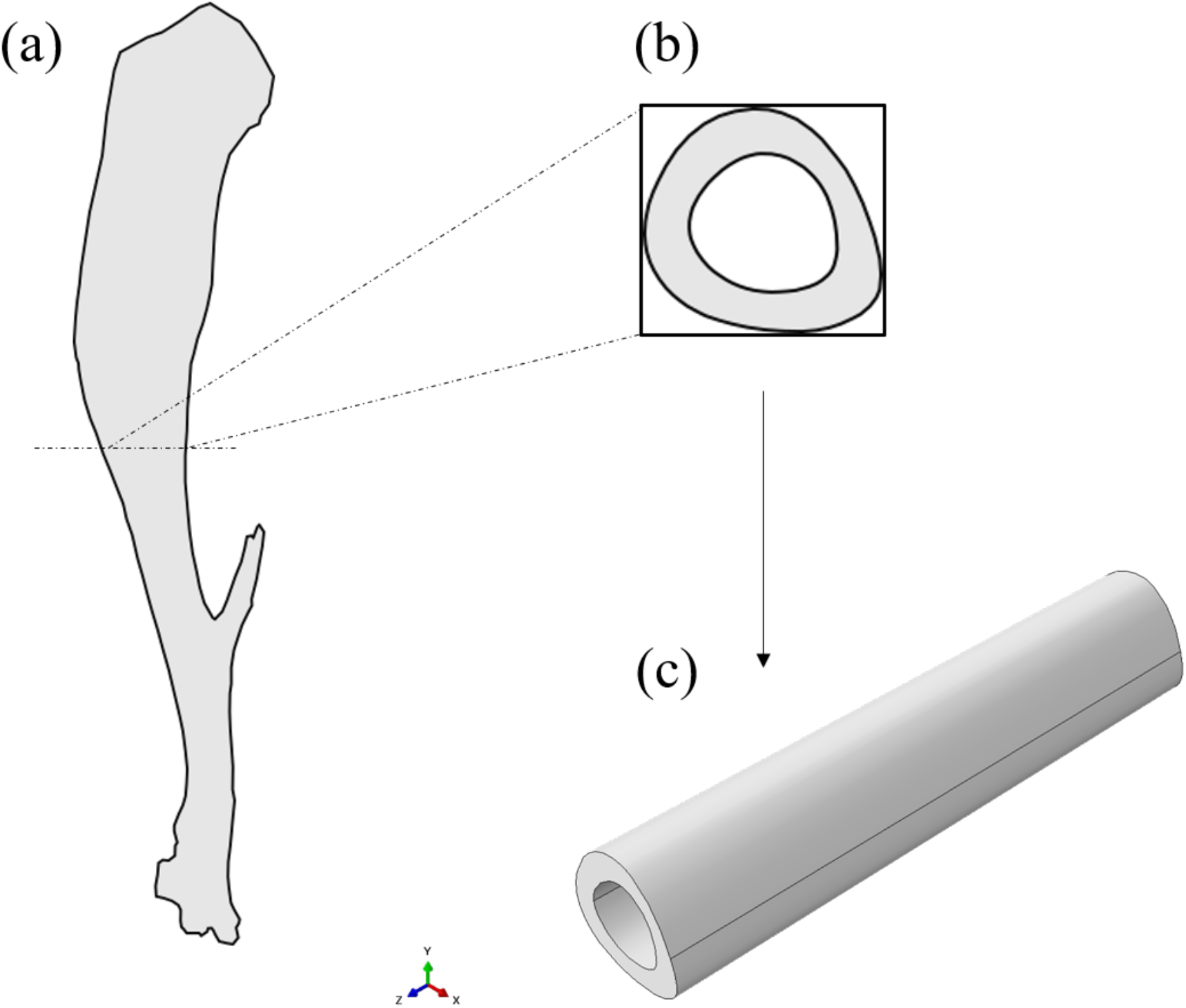
Pictorial representation of (a) 8-week-old C57BL/6 mouse tibia with (b) an idealized cross section at section a-a’ (adapted from Barmen et al. [22]). (c) A simplified beam developed with a cross-section similar to the section at a-a’.

#### 2.1.2 Porosity and Permeability of Bone

The cortical bone displays distinct porosities across scales, including vascular porosity, lacune canalicular porosity, and collagen apatite porosity [23]. We focused on lacunar-canalicular porosity only as it is believed to play a crucial role in bone adaptation due to osteocytes in lacunae and their processes in canaliculi [2].

Even though bone intrinsic permeability has been extensively explored in the last decade, the computational and experimental estimation of intrinsic permeability ranges between 10^−18^ – 10^−22^ m^2^ and 10^−22^ – 10^−25^ m^2^, respectively [2, 24-27]. Experimentally measuring the relaxation time provides a means of determining the permeability value [26, 28]. Pereira et al. [6] estimated the intrinsic permeability of the order of 10^−22^ m^2^ for the previously estimated relaxation time of 6.76 sec by Zhou et al. [10]. Accordingly, for this simulation, the permeability of the bone was considered to be 3x10^−22^ m^2^.

#### 2.1.3 Permeability of Bone Surface

There is no consensus about the permeability of the periosteum. Evans et al. [8] have experimentally determined the permeability of the periosteum to be 10^−17^ m^2^ ex-situ and thus suggested that the periosteum to be highly permeable. Carriero et al. [7] have used the permeability measured by Evans and shown that periosteal permeability of an order of 10^−17^ m^2^ (which is actually more than the permeability of the bone) significantly overestimates the periosteal adaptation. In contrast, Li et al. [29] have indicated that the periosteum (external bone surface) is impermeable. Steck et al. [30] characterized the periosteum as relatively impervious and the endosteum (inner surface of the bone) as relatively permeable. Price et al. [9] also experimentally measured the high circumferential fluid velocity in situ, which according to Zhou et al. [10] indicates the periosteum’s very low (negligible) permeability. Given the above studies, we preferred Price et al.’s in-situ findings and accordingly assumed the permeability of the periosteum negligible. Endocortical surfaces, on the other hand, are considered fully permeable in accordance with the literature.

#### 2.1.4 Loading and Boundary Conditions

To predict the site-specific new bone formation at both cortical surfaces, we modeled the in-vivo study by Barmen et al. [22]. The loading was referred to as mid strain regimen, where 8-week-old C56BL/6J mice were subjected to axial loading of 8.8 N, engendering 2050 µƐ on the anteromedial surface of the midsection. The loading profile consists of 4 haversine waveforms at a frequency of 2 Hz followed by a rest of 3 seconds at the maximum load and repeated the loading profile 55 times a day. The loading was applied for 14 days, with three days of loading followed by a rest of one day. We simulated the above loading protocol by setting zero displacements to nodes at one end of the beam and eccentric compressive loading at the other to induce the strains on the beam’s mid-cross-section, similar to tibial midsection strains in the experimental setup. The load was applied to create a maximum axial strain of 2050 µƐ at the anteromedial surface and a posterior-to-anterior axial strain ratio of 1.5-2 [31].

The boundary conditions of impermeable and permeable periosteal and endosteal surfaces, respectively, were implemented in the model by setting the zero pressure, i.e., (***P*** = 0), and zero flow, i.e., (∇***P***. *r* = 0) at the endocortical and periosteal surface, respectively.

#### 2.1.5 Other Details

After conducting the convergence study, as shown in Fig. 3, the beam was meshed with 81600 C3D8RP (Continuous 3 Dimensional 8-noded Reduced-integration Pore-pressure hexahedral) elements with an average distance of 50 µm between the nodes. A minimum increment step time of 2x10^−3^ sec was used for guaranteed convergence. The material properties presented in Table 1 were taken directly from Zang and Cowin [32] and Cowin [23], except for the permeability of the bone.

**Table 1.** Material properties used for simulation (adapted from [32] and [23])

| Property | Bone | Units |
| --- | --- | --- |
| Young Modulus (E) | 12000 | MPa |
| Drained Poisson Ratio ($\nu$) | 0.3 | |
| Bulk Modulus Solid ($K_s$) | 17000 | MPa |
| Bulk Modulus Liquid ($K_f$) | 2300 | MPa |
| Porosity ($n_p$) | 5% | |
| Intrinsic Permeability | $3 \times 10^{-21}$ | $m^2$ |
| Specific Weight ($\gamma$) | $9.8 \times 10^3$ | $Nm^{-3}$ |
| Dynamic Viscosity ($\mu$) | $8.9 \times 10^{-2}$ | Pa s |

**Fig. 3.**
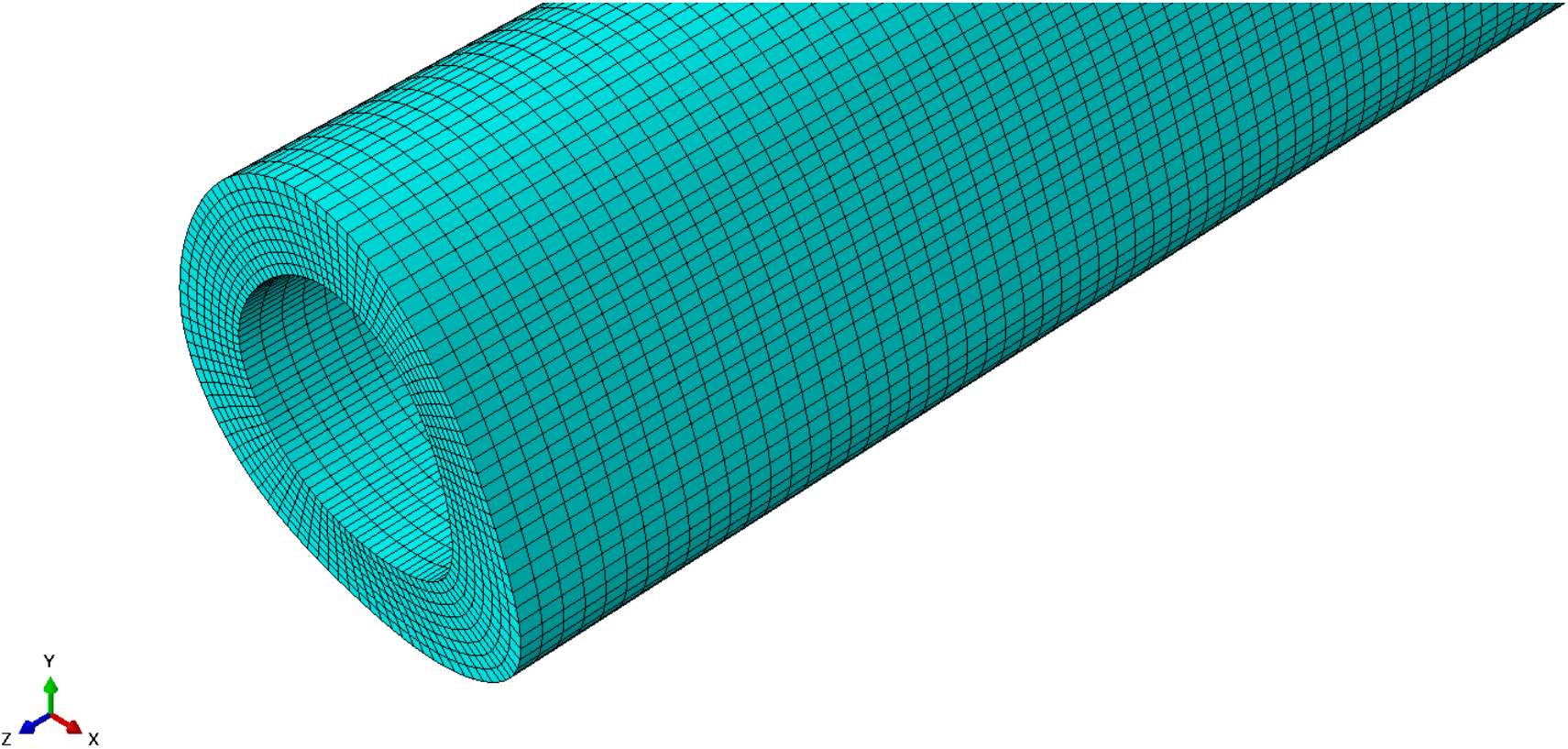
Finite element meshed beam having a cross-section similar to the midsection of mouse tibia.

### 2.2 Dissipation energy density

#### 2.2.1 Dissipation Energy at the Cellular Level

As discussed in the Introduction Section, there are three cellular stimuli: pore pressure, tissue strain, and fluid flow. The energy dissipated at the tissue level under the action of the load can be considered as a combination of energy dissipation due to (i) viscous fluid flow, (ii) pore pressure, and (iii) strain exerted on the osteocytes. Hence, dissipation energy may be formulated as follows:

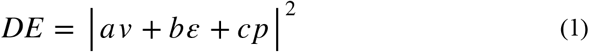

where, *ν ε*, and *p* are the fluid velocity, tissue-level volumetric strain, and pore pressure.*a, b, c* and are the constants. |x| denotes the amplitude of the quantity x.

However, the mechanical strain needed to activate the bone cells in culture is 10-100 times larger than in vivo bone strain [15]. Hence, the effect of tissue-level strain on osteocytes is minimal and can be neglected. Thus, Eq. (1) may be rewritten as:

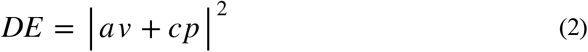

As the periosteal and endocortical surfaces are assumed to be ideally impermeable and permeable, respectively, the fluid velocity is assumed to be negligible at the periosteal surface, whereas pressure is assumed to be zero at the endocortical surface. Accordingly, the final equation is the following:

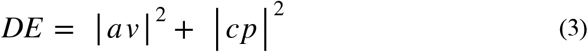

The above relation is also approximately true between endocortical and periosteal surfaces, as the pore pressure and fluid velocity are approximately orthogonal to each other because of the boundary conditions and the relation between them. The variation of pressure along the cortical thickness may be assumed to be approximately varying sinusoidally with a node at the endocortical surface and a peak at the periosteal surface, in accordance with the literature [32-34]. The fluid velocity (e.g., a cosine function) is proportional to the gradient of the pore pressure (e.g., a sine function), and the resultant of the two (*a v* and *c p*) can be obtained as the square root of the summation of the squares of amplitudes of the two (*a v* and *c p*), in accordance with Eq. (3).

Thus, the tissue-level total dissipation energy density will be a combination of dissipation energy density due to fluid flow and that due to pore pressure.

#### 2.2.2 Dissipation energy density – due to fluid flow alone

The energy dissipated due to the viscous nature of fluid over a loading cycle time period (*T*) can be considered a stimulus and is defined as [5]

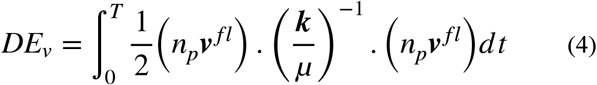

where *DE*_*v*_ is the dissipation energy density per cycle due to fluid flow, *n*_*p*_ is the porosity, ***v*** ^*f l*^ is the fluid velocity, ***k*** is the intrinsic permeability of cortical bone tissue, and of the kinematic viscosity of the fluid in LCN.

#### 2.2.3 Dissipation energy density – due to pressure alone

Osteocytes can be modeled as a viscoelastic material as it shows time-dependent behavior [35, 36]. In viscoelastic material, the hysteretic loss (dissipation energy density for one cycle) can be expressed as

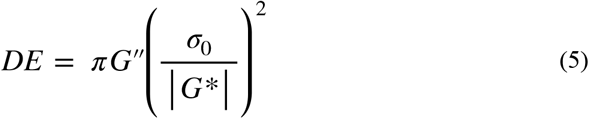

where *G\** = *G*′ + *iG*′′ and σ_0_ are the complex modulus and stress amplitude, respectively [37].

Assuming stress amplitude induced inside an osteocyte (σ _*o*_) to be directly proportional to pressure amplitude (*p*_0_) acting on the surface of the osteocyte, i.e., can be estimated as *σ*_*o*_ ∝ *p*_0_, dissipation energy density due to pressure acting on osteocytes can be estimated as

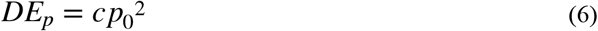

where *c* is the proportionality constant that needs to be determined.

#### 2.2.4 Zone of Influence

Osteocytes form the network structure in bone by connecting their processes via gap junction [38]. The processes of osteocytes also form the gap junction with osteoblasts/lining cells on the bone surface. It suggests that the osteocytes in the LCN detect the mechanical cues, convert them into biochemical signals, and transmit them to the osteoblasts on the periosteal and endocortical surfaces through diffusion [39]. This communication is implemented by Prasad and Goyal as diffusion [40]. However, it has been modeled differently in the existing models, where a spherical zone of influence is considered instead [5, 41, 42]. We considered the zone of influence for its simplicity, which suggests that osteocytes closer to the surface contribute more than the osteocytes away from the surface. Averaging was weighted with the exponential function *w*|*x*|.

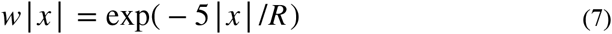

where *| x|* is the distance between the node of interest at the surface and the inner node in the zone of influence.

The value of ***R*** was set equal to the radius of an osteon, i.e., 150*μm* [43]. The modified stimulus due to fluid flow and pressure at node *i* is calculated as a weighted average in the zone of influence as

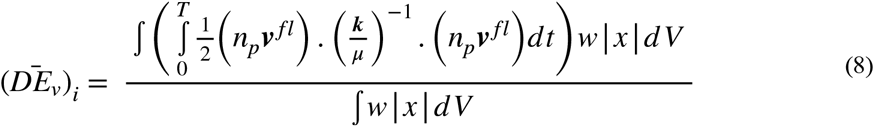

or

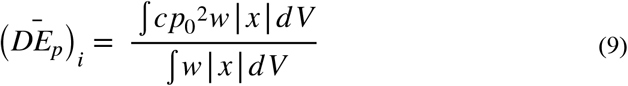

where *V* is the volume of the zone of influence.

### 2.3 Derivation of Mineral Apposition Rate (MAR)

When an exogenous harmonic force is applied to the bone, the overall response to increase bone cross-sectional area (*A*_*c*_) (for example, per day) may be assumed to be similar to that of a first-order control system, i.e.

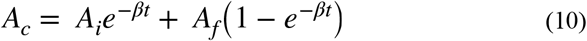

or

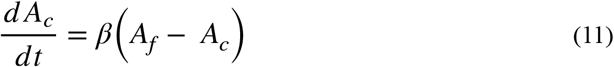

where *A*_*c*_, *A*_*i*_, and *A*_*f*_ are the current, initial, and final cross-sectional areas of the bone, respectively. *β* is an unknown function of loading parameters such as the number of load cycles *N* etc. (to be determined), and *t* is the time. This comparison is drawn because bone adaptation responds to the stimulus (energy dissipation) from the current week’s loading with a diminishing return compared to the previous week’s load. This reduced response is due to the increased cross-sectional area, which enhances the stiffness of the bone. As a result, bone adaptation gradually reaches saturation over time. Thus, we adopted the exponential saturation in bone adaptation.

It is desired to express *d A*_*c*_ /*dt* (i.e., the rate of change in bone cross-sectional area with respect to time) in terms of dissipation energy density. The dissipation of energy at the tissue level in bone is a combination of energy dissipated due to fluid flow, pressure, and strain at the cellular level. The tissue-level behavior of dissipation energy density is the same whether we consider bone viscoelastic or poroelastic materials. Therefore, dissipation energy density due to viscoelasticity or poroelasticity can be used interchangeably at the tissue level. Hence for the sake of simplicity, we used viscoelastic dissipation energy density in the derivation. The tissue-level dissipation energy density per cycle of a sinusoidal loading of amplitude *F*_0_, is as follows:

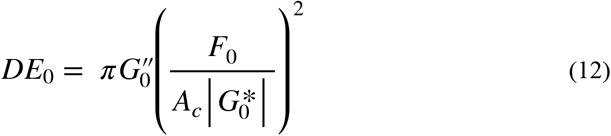

where 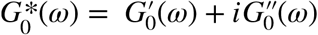 is the complex modulus (as a function of forcing frequency) for the bone tissue corresponding to in-vivo conditions. 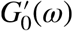 and 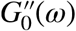 respectively are the corresponding storage and loss moduli.

If there are *N* cycles of the load being applied per day for *d* number of days per week, the total dissipation energy density may be given by

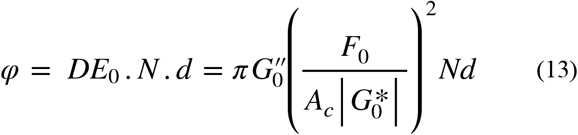

or

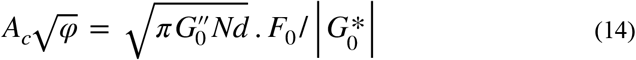

If the same loading protocol is maintained for the same bone, then for the bone cross-section in consideration:

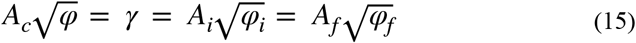

where *γ* is a constant for a given loading protocol. Subscript *i* refers to the ‘initial condition’, i.e., the time immediately after the exogenous loading on bone starts. Subscript *f* refers to the ‘final condition’, i.e., the time by which the bone cross-section is completely adapted to the exogenous loading. Equation (15) may be applied to Eq. (11) to obtain the following:

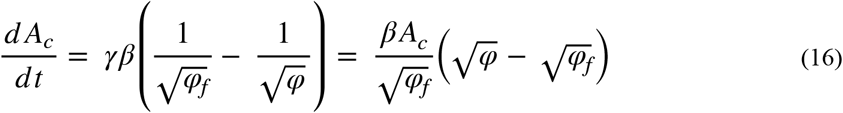

In experimental studies, the site-specific Mineral Apposition Rate (MAR) is approximately calculated for ‘initial conditions’ (i.e., within 2-3 weeks of initiation of exogenous loading). Hence Eq. (16) may be rewritten as:

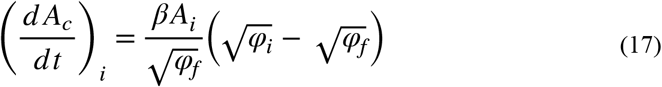

The whole bone may be divided into many sectors (Fig. 4), the area of each of which may be used to calculate local MAR (i.e., new bone thickness per unit time) as follows:

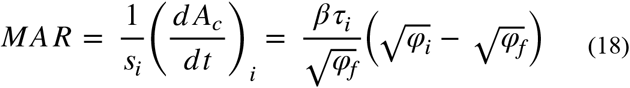

where *s*_*i*_ is the circumferential length for the sector in consideration and *A*_*i*_ may now be considered as the sector area. The ratio of *A*_*i*_ and *s*_*i*_ is the cortical thickness (*τ*_*i*_) for the sector. The simplified approximate version of Eq. (18) may be given by:

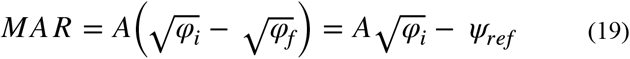

where *A* is a constant. *φ*_*f*_ may also be termed a universal threshold value. 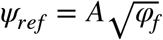 is again a function of *N* and *d*. The function *β* is given below:

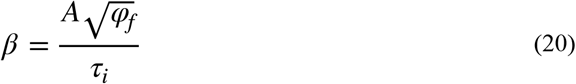

**Fig. 4.**
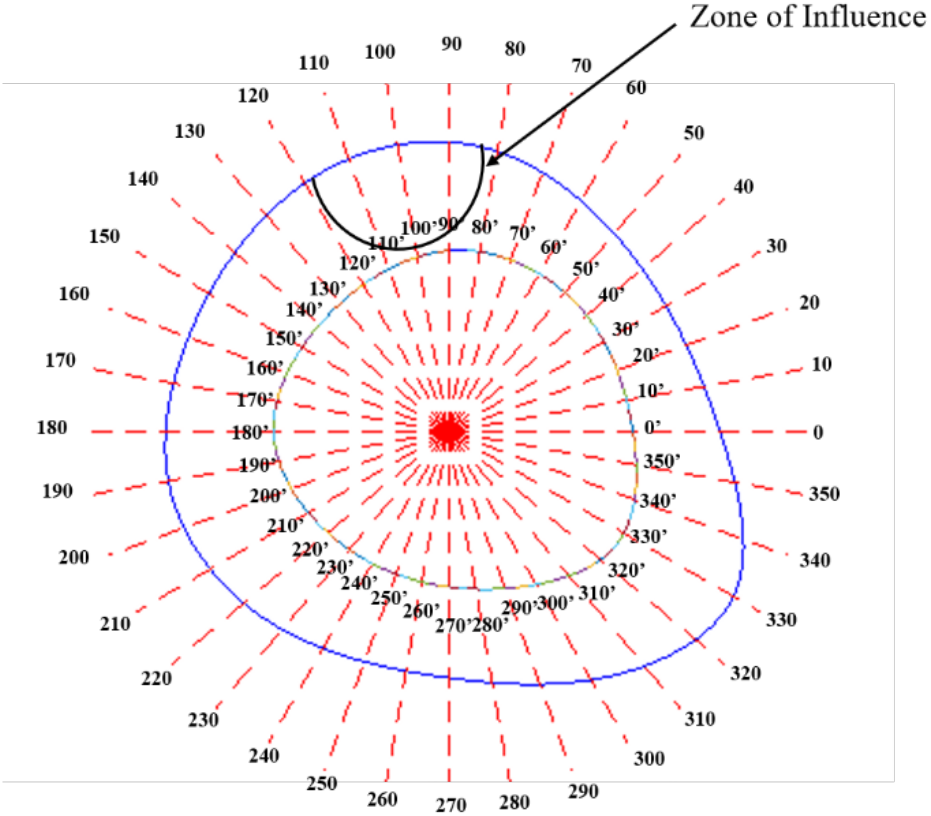
Cross-section of the mid diaphysis of 8-week-old C57Bl/6J mice tibia showing (a) zone of influence and (b) radial lines (adapted from Fig 4(b) of Berman et al. [22]).

### 2.4 Osteogenesis as a function of fluid flow

To measure the site-specific new bone formation, we divided the circumference of the mid-cross section of the beam into 360 points by the radial lines passing through the centroid of the cross-section. The lines intersecting the periosteal and the endocortical rim were numbered *i* = 1 – 360 and *i* = 0’ – 360’ in an anti-clockwise direction, respectively. For clarity, these lines are shown at an angle of ten degrees in Fig. 4. At these intersection points at periosteal and endosteal surfaces; we estimated the site-specific osteogenesis. Following Eq. (19), we hypothesized that the quantity that predicts the spatial new bone thickness per unit time on the considered section and surface (endocortical and periosteal) is given by

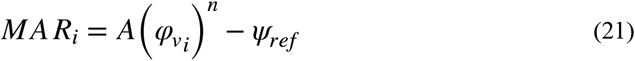

where 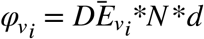 and 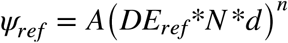. *MAR*_*i*_ is the mineral apposition rate at a given node and 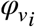 is the corresponding stimulus due to fluid flow. *A* is the remodeling rate to be determined, *N* is the number of loading cycles, and *d* is the number of loading days. The power of *n* is another constant to be determined, which according to Eq. (19), is expected to be around 0.5. *ψ*_*ref*_ may be considered as the reference MAR existing in the absence of exogenous mechanical loading. *DE*_*ref*_ is the threshold/reference value of dissipation energy density for one cycle, and 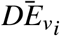 which is actual dissipation energy due to external loading, must exceed it for new bone formation to occur. Equation (21) can fit the MAR data for the endocortical surface as follows.

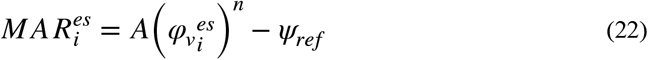

Where *e s* represents the endocortical surface.

The unknown model parameters, i.e.,*A, n* and *ψ*_*ref*_ can be estimated by minimizing the error squared between the predicted mineral apposition rate 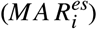and the corresponding experimental value 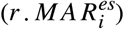at nodes *i* on the endocortical surface using Eq. (23).

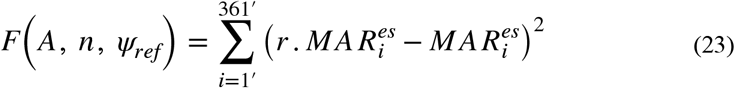

The value of 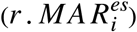 which signifies the relative MAR is calculated by subtracting the natural bone formation data of contralateral bones from the loaded bone to see the loading effect.

### 2.5 Osteogenesis as a function of pore pressure

Several in vitro and in vivo studies have shown that hydrostatic pressure is essential in bone maintenance [44, 45]. Hence, similar to the previous section, a site-specific model is developed to predict the osteogenesis at the cortical surfaces; however, this time with pore pressure as a stimulus. The quantity that estimated the new bone distribution as a function of fluid pore pressure is given by:

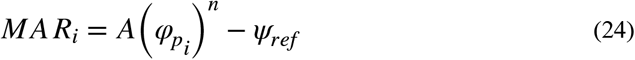

where 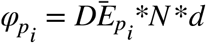 and 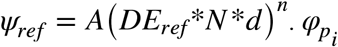 is the total dissipation energy density due to pore pressure at a given node *i* and 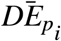 is a dissipation energy density due to pore pressure during one cycle of external loading.

As the pore pressure is assumed to be zero at the endocortical surface (which will result in a negligible dissipation energy density), the values of the parameter, *A, n* and *ψ*_*ref*_ are estimated by fitting the new bone distribution at the periosteal surface as:

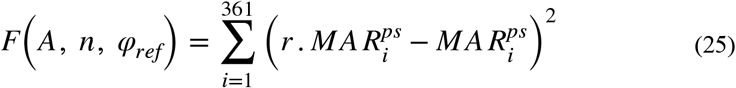

where *p s* represents the periosteal surface. 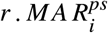 and 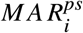 are the experimental and computationally estimated site-specific MAR at a given node *i*.

### 2.6 Osteogenesis as a function of fluid flow and pore pressure

We use the same in vivo data as in subsection 2.4 to estimate the site-specific new bone thickness at cortical surfaces as a function of fluid flow and pore pressure. The quantity that predicts the site-specific new bone formation at both surfaces is given by Eq. (26).

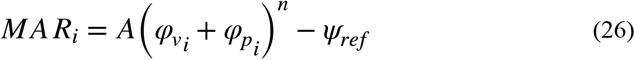

where 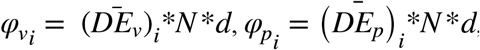 and *ψ*_*ref*_ = *A*(*DE*_*ref*_ * *N*\**d*)^*n*^.

Due to the different permeability of the cortical envelope, the fluid flow and fluid pore pressure will dominate on the endocortical and the periosteal surface, respectively. Therefore, the value of parameters, i.e., *A,c, n*, and *φ*_*ref*_ are estimated by minimizing the error squared between the predicted mineral apposition rate (*MAR*_*i*_) and the corresponding experimental value (*r p*. *MAR*_*i*_) of the periosteal and endocortical surfaces using Eq. (27).

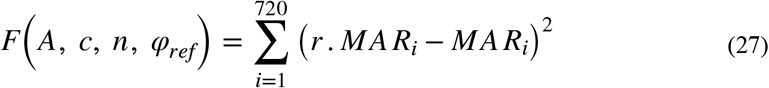

### 2.7 Statistical Analysis

The statistical analyses are the same as that adopted by Prasad and Goyal [40]. To compare the numerical predictions to the experimental data, the total new bone area formed per unit of time is obtained by integrating the MAR (i.e., new bone thickness formed per unit of time) over the circumference of each of the two bone surfaces at the cross-section under study. The bone formation rate per unit bone surface (BFR/BS), or simply, bone formation rate (BFR), is obtained by dividing the new bone area formation rate by the total perimeter of the surface in consideration. Then the Student’s t-test (a one-sample, two-tailed t-test) was used to compare the bone formation rate (BFR) predicted by the mathematical model to the corresponding experimental value. The mineral apposition rate (MAR) is recorded on a circular scale, and therefore, Watson’s U^2^ test is used to compare the experimental and simulated MAR [46, 47]. The Student’s t-test and Watson’s U^2^ test have been done using MATLAB (MathWorks Inc.) programming.

## 3 Results

The model simulates osteogenesis as a function of fluid flow alone, pore pressure alone, and fluid flow with pore pressure at cortical surfaces. The computed strain distribution (Fig. 5a) for mid strain loading protocol of Barmen et al. [22] shows that the maximum strain induced at the anteromedial surface is 2056 µƐ, which is close to the experimental strain of 2050 µƐ. The velocity is negligible at the periosteal surface, which reaches the maximum at the endocortical surface (see Fig. 5b). In contrast, pressure is zero at the endocortical surface and reaches the maximum at the periosteal surface (see Fig. 5c). We estimated the dissipation energy density (*DE*_*v*_ and *DE*_*p*_) for one cycle as given by Eq. (4) for fluid flow and Eq. (6) for fluid pressure. Then, the results were extrapolated according to the loading cycles and number of days in the experimental protocols to calculate the total dissipation energy density (*φ*_*i*_). Figure 6 shows the plot of dissipation energy density due to fluid flow and pore pressure along the thickness of the beam mid-cross section. The dissipation energy density due to fluid flow follows the trend of fluid velocity, i.e., it is maximum at the endocortical surface and reaches its minimum value at the periosteal surface. Similarly, dissipation energy density due to pore pressure also follows the trend of pore pressure, i.e., it is minimum at the endocortical surface and maximum at the periosteal surface.

**Fig. 5.**
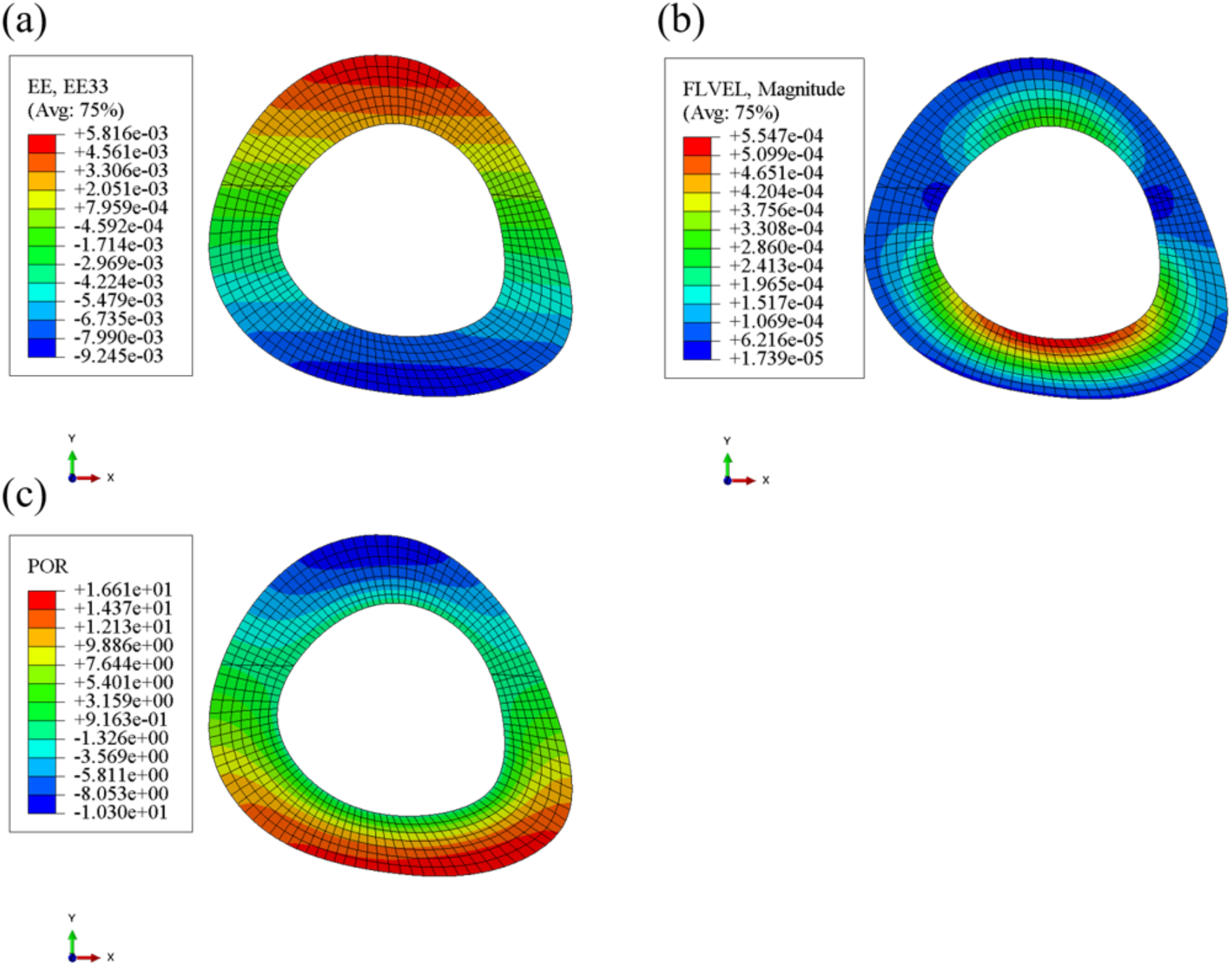
(a) Longitudinal strain distribution adapted from Barmen et al. [22], (b) velocity distribution (mm/sec), and (c) pressure distribution (MPa).

**Fig. 6.**
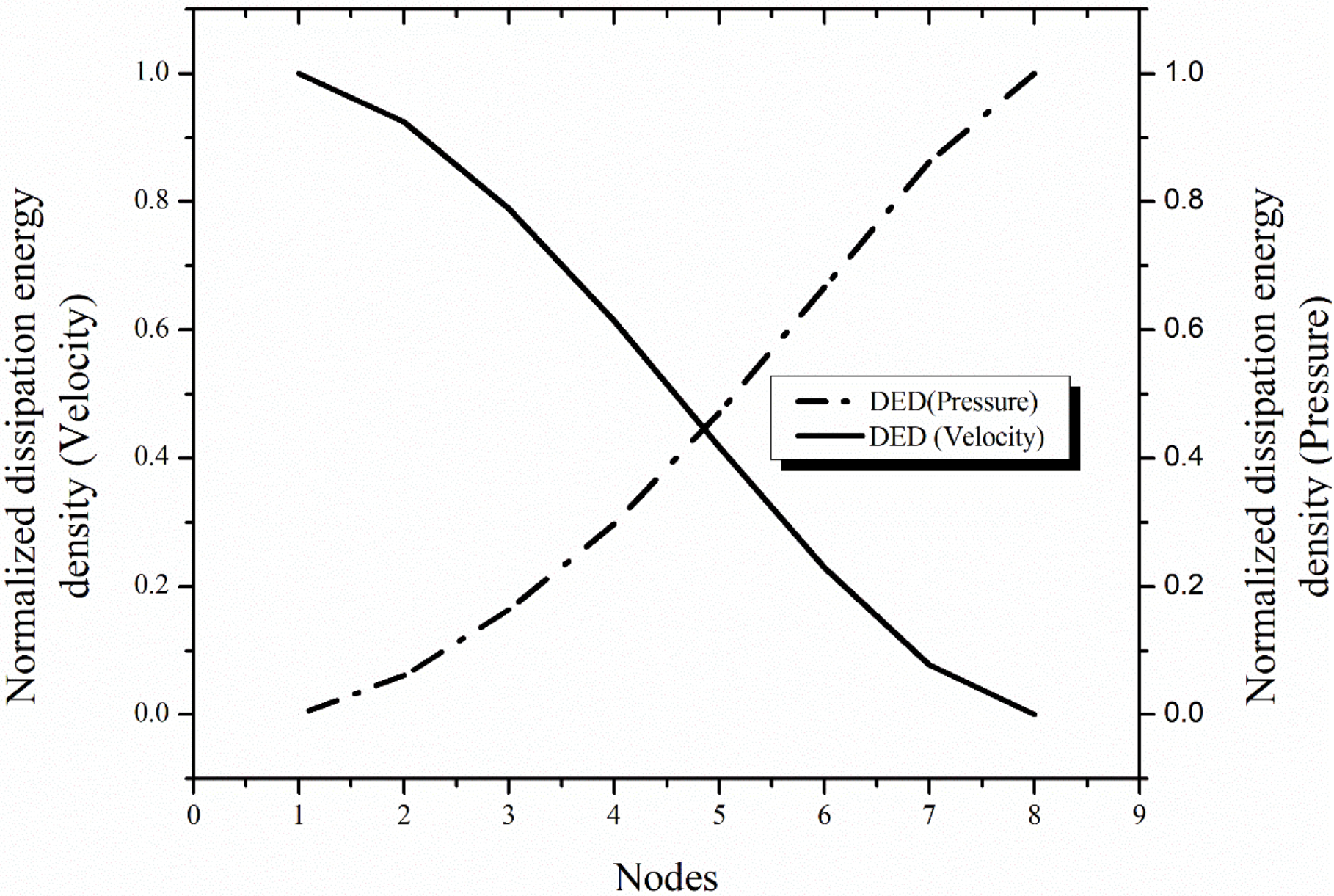
Dissipation energy density (DED) due to fluid velocity and pore pressure along the radial line.

### 3.1 Osteogenesis as a function of fluid velocity

The potential of fluid flow to predict osteogenesis at both cortical surfaces has been investigated here. The values of parameters that fit the observed new bone thickness at the endocortical surface of Barmen et al. [22] and relate to the dissipation energy density only due to fluid flow are as follows: *A* = 184.94 µm/√N/day, *n* = 0.401 and *ψ*_*ref*_ = 0.374 µm^3^/µm^2^/day. Corresponding to the computed *ψ*_*ref*_, *DE*_*ref*_ comes out to be 3.862x10^−10^ Nµm/µm^3^. Similar to the in vivo new bone distribution at the endocortical surface (Fig. 7a), the model also predicted the site-specific new bone formation at the anterior and posterior sides of the endocortical surface (Fig. 7b). The BFR calculated by the model is 0.6316 µm^3^/µm^2^/day at the endocortical surface, which is not significantly different from the experimental BFR of 0.5964 ± 0.14285 µm^3^/µm^2^/day (*p*-value = 0.812, t-test). Additionally, the statistical significance of site-specific MAR at the endocortical surface is measured using Watson’s U^2^ test, which shows that the computed MAR is not significantly different from the experimental MAR (*p*-value = 0.5957, Watson’s U^2^ test). However, the predicted BFR of 0.0694 µm^3^/µm^2^/day at the periosteal surface is significantly different from the experimental BFR of 0.9257 ± 0.15855 µm^3^/µm^2^/day (*p*-value = 0.000432, t-test). The site-specific new bone distribution at the periosteal surface Fig. 7(b) is also significantly different from the experimental MAR (*p*-value = 0.001509, Watson’s U^2^ test).

**Fig. 7.**
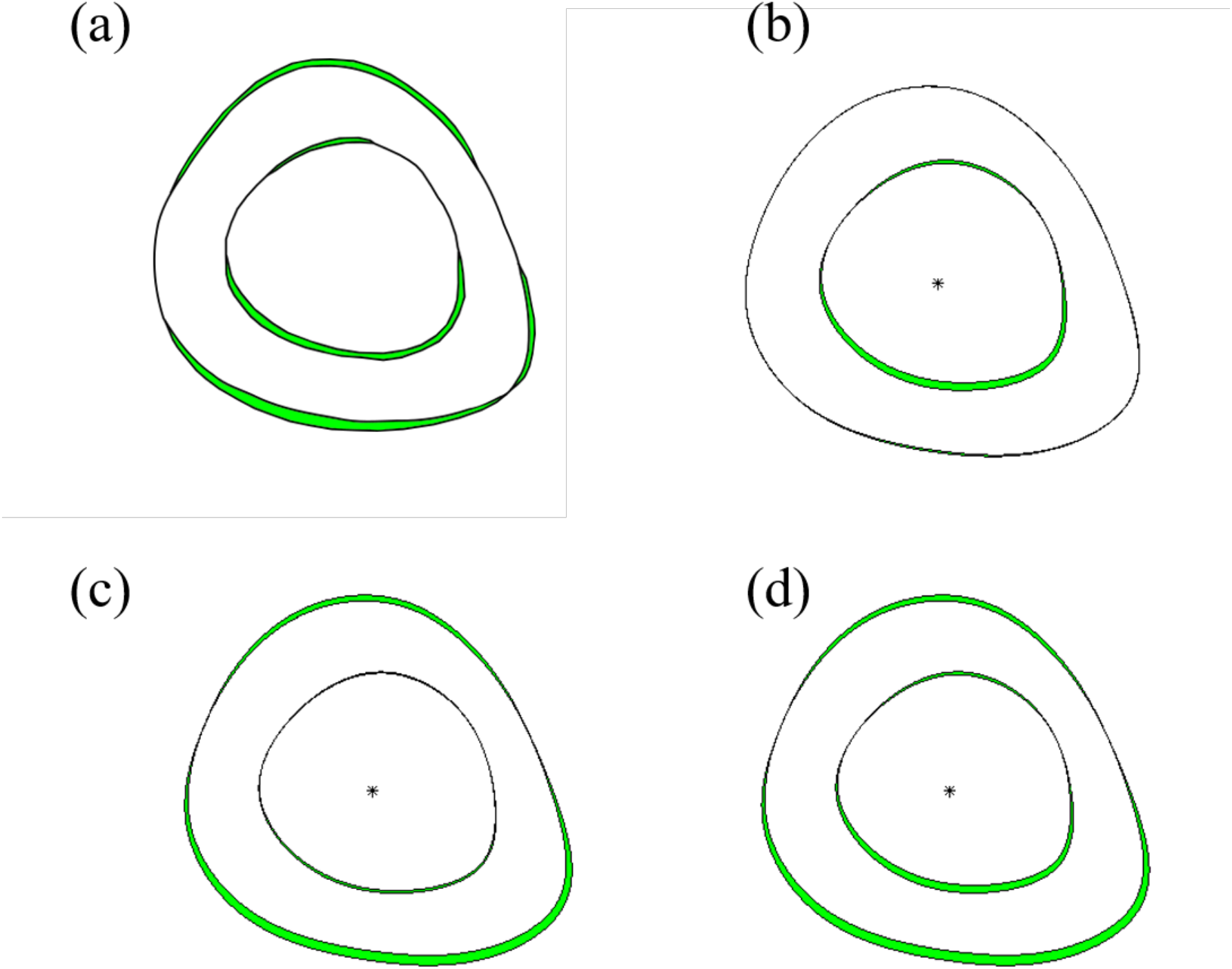
(a) In vivo, new bone formation at the cortical surfacers for mid-strain loading protocol (adapted from Berman et al. [22]), and the corresponding numerically predicted new bone distribution as a function of (b) fluid flow only, (c) pore pressure only, and (d) fluid flow with pore pressure.

### 3.2 Site-Specific model for pore pressure

The osteogenic potential of pore pressure alone has been investigated here. The value of parameters that fit the MAR data on the periosteal surface come out to be as follows: *Ac*^*n*^ = 2.06, = 0.5196, and *ψ*_*ref*_ = 0.0765 µm^3^/µm^2^/day. *c* cannot be computed separately; hence, *Ac*^*n*^ is determined. The value of *DE*_*ref*_ is 3.5624x10^−6^ Nµm/µm^3^. Pore pressure as a stimulus in the model predicts the new bone formation at the periosteal surface (Fig. 7c). However, it fails to model osteogenesis completely at the endocortical surface. The BFR/BS estimated at the periosteal and endocortical surface is 0.9872 µm^3^/µm^2^/day (*p*-value = 0.7071, t-test) and 0.219 µm^3^/µm^2^/day (*p*-value = 0.0268, t-test). The corresponding experimental values are 0.9257 ± 0.15855 µm^3^/µm^2^/day and 0.5964 ± 0.14285 µm^3^/µm^2^/ day. Site-specific distribution at the periosteal surface, as shown in Fig. 7c, is not significantly different from the experimental distribution (*p*-value = 0.8685, Watson’s U^2^ test). However, the new bone distribution at the endocortical surface does not come close to the in vivo bone distribution (*p*-value = 0.0381, Watson’s U^2^ test).

### 3.3 Site-Specific model for fluid velocity and pore pressure

The value of the parameters *A,c,n* and, *φ*_*ref*_ which fit the new bone thickness and dissipation energy density due to both fluid flow and pore pressure are as follows: *A* = 503.94 µm/√N/day,*c* = 1.8534x10^−5^,*n* = 0.497, and *ψ*_*ref*_ = 0.252 µm^3^/µm^2^/day. *DE*_*ref*_ is estimated to be 4.61x10^−10^ Nµm/µm^3^. The osteogenesis results at the cortical surface improve when fluid flow and fluid pressure are considered together (see Fig. 7d). The computed BFR/BS of 0.9709 µm^3^/µm^2^/day and 0.6276 µm^3^/µm^2^/day at the periosteal and endocortical surface is not significantly different from the experimental BFR of 0.9257 ± 0.15855 µm^3^/µm^2^/day (*p*-value = 0.782, t-test) and 0.5964 ± 0.14285 µm^3^/µm^2^/ day (*p*-value = 0.832, t-test) at the corresponding surfaces. The distribution of newly formed bone in the experimental study and predicted by the mathematical model at both surfaces are also not significantly different (*p*-value = 0.9465 at the periosteal surface and *p*-value = 0.714 at the endocortical surface, Watson’s U^2^ test) (Fig. 7d).

### 3.4 Simplified Model

In contemporary mathematical models, bone adaptation is commonly presumed to be proportional to either strain energy density or dissipation energy density [41, 48, 49]. However, this foundational assumption may be misguided, as in vivo research has revealed that bone adaptation evinces a linear dose-response association with strain [13]. This implies that bone adaptation may be more appropriately modeled as being proportional to the square root of dissipation energy density [50]. This is supported by the derivation of MAR in section 2.3, where MAR is shown to vary with the square root of dissipation energy density. In the original model with unconstrained *n* (see section 3.3) too, the value of exponent *n* comes out to be 0.497, and thus may act as another validation for MAR to be approximately proportional to the square root of dissipation energy density. Therefore, for the sake of simplicity and in accordance with the derived Eq. (19), we assumed the value of *n* to be 0.5 and tested the hypothesis, whether the site-specific mineral apposition rate at both periosteal and endocortical surface is directly proportional to the square root of dissipation energy density (above its reference value), i.e.,

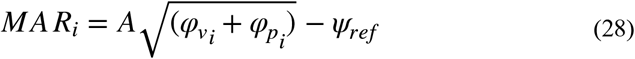

where 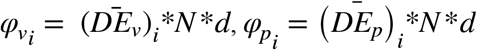 and 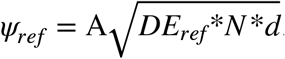.

Similar to section 3.3, when *n* = 0.5, the value of parameters, i.e.*A*, and *ψ*_*ref*_ come out to be 509.33 µm/√N/day and 0.227 µm^3^/µm^2^/sec, respectively. The value of *DE*_*ref*_ calculated to be 3.889x10^−10^ N-µm/µm^3^. The BFR estimated at the periosteal surface by the simplified model is 0.9658 µm^3^/µm^2^/sec, which is significantly similar to the BFR of 0.9257 ± 0.15855 µm^3^/µm^2^/sec obtained experimentally (*p*-value = 0.806, t-test). According to the Watson U^2^ test, the simplified model’s estimation of new bone distribution at the periosteal surface, as depicted in Fig. 8b, is not significantly distinct from the actual new bone distribution in vivo, as shown in Fig. 8a (*p*-value = 0.9522, Watson U^2^-test). Similarly, the BFR computed on the endocortical surface by the simplified model is 0.6291 µm^3^/µm^2^/sec (Fig. 8b), which is not significantly different from the in vivo BFR of 0.5964 ± 0.14285 µm^3^/µm^2^/sec (*p*-value = 0.8241, t-test). The Watson U^2^ test also shows that simulated bone distribution at the endocortical surface (Fig. 8b) is not significantly different from the in vivo new bone distribution, as shown in Fig. 8a (*p*-value = 0.6654, Watson U^2^-test).

**Fig. 8.**
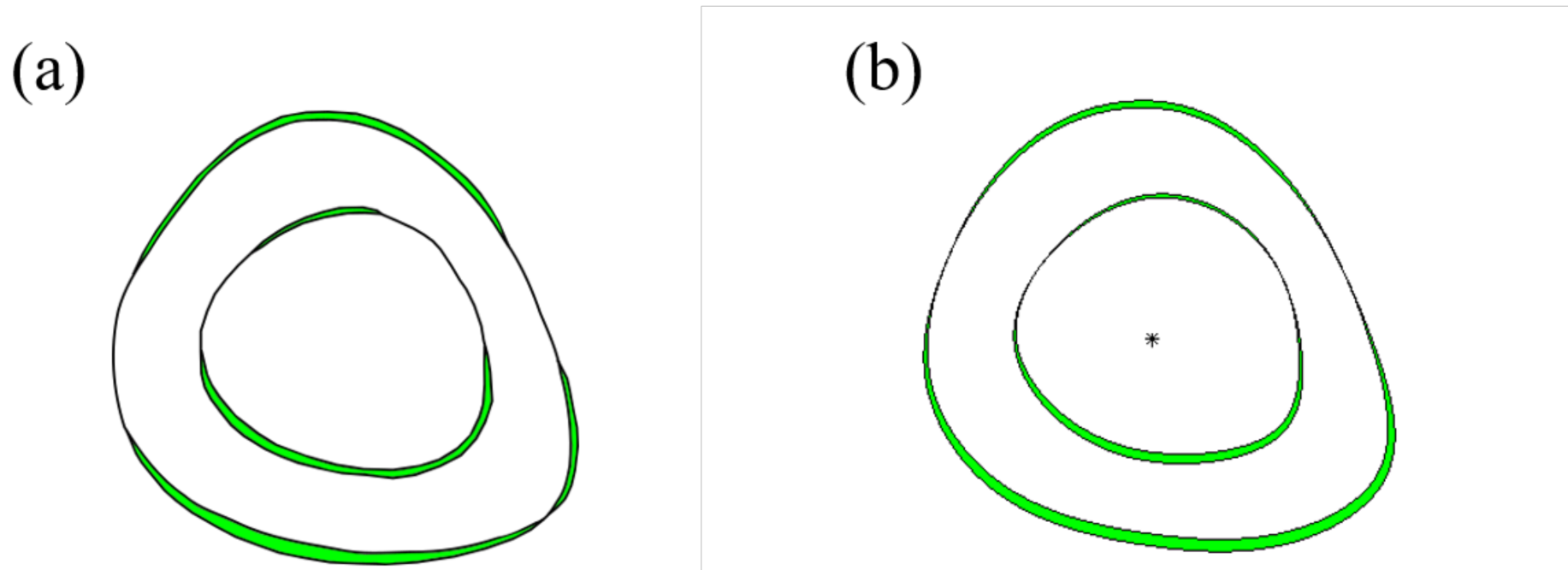
(a) The distribution of new bone in vivo at cortical surfaces for mid-strain loading protocol (adapted from Berman et al. [22]). (b) The corresponding computationally estimated new bone distribution using a simplified model is also illustrated.

#### 3.4.1 Prediction Example

To verify the robustness of the model, we solved a high-strain loading example taken from Berman et al. [22]. The difference is that instead of 2050 µƐ, 2400 µƐ is applied at the anteromedial site. The value of parameters *A, c, n*, and *DE*_*ref*_ remain the same as that in section 3.4. The predicted BFR at the periosteal surface is 1.5618 µm^3^/µm^2^/ sec, which is close to the experimental BFR of 1.672 ± 0.1457 µm^3^/µm^2^/sec (p-value = 0.491, t-test). Similarly, corresponding to the experimental BFR of 1.0143 ± 0.105 µm^3^/µm^2^/sec at the endocortical surface, the BFR measured from the model is 1.0205 µm^3^/µm^2^/sec, which is close to the experimental BFR (p-value = 0.955, t-test). The new bone distribution at both the cortical surfaces, i.e., MAR (Fig.9b), is not significantly different from the experimental new bone formation at the mid-diaphysis (Fig. 9a) (p-value = 0.441 and p-value = 0.08 for periosteal and endosteal surfaces, Watson’s U^2^ test). However, the model underestimates the new bone distribution at the anterior side of the endocortical surface.

**Fig. 9.**
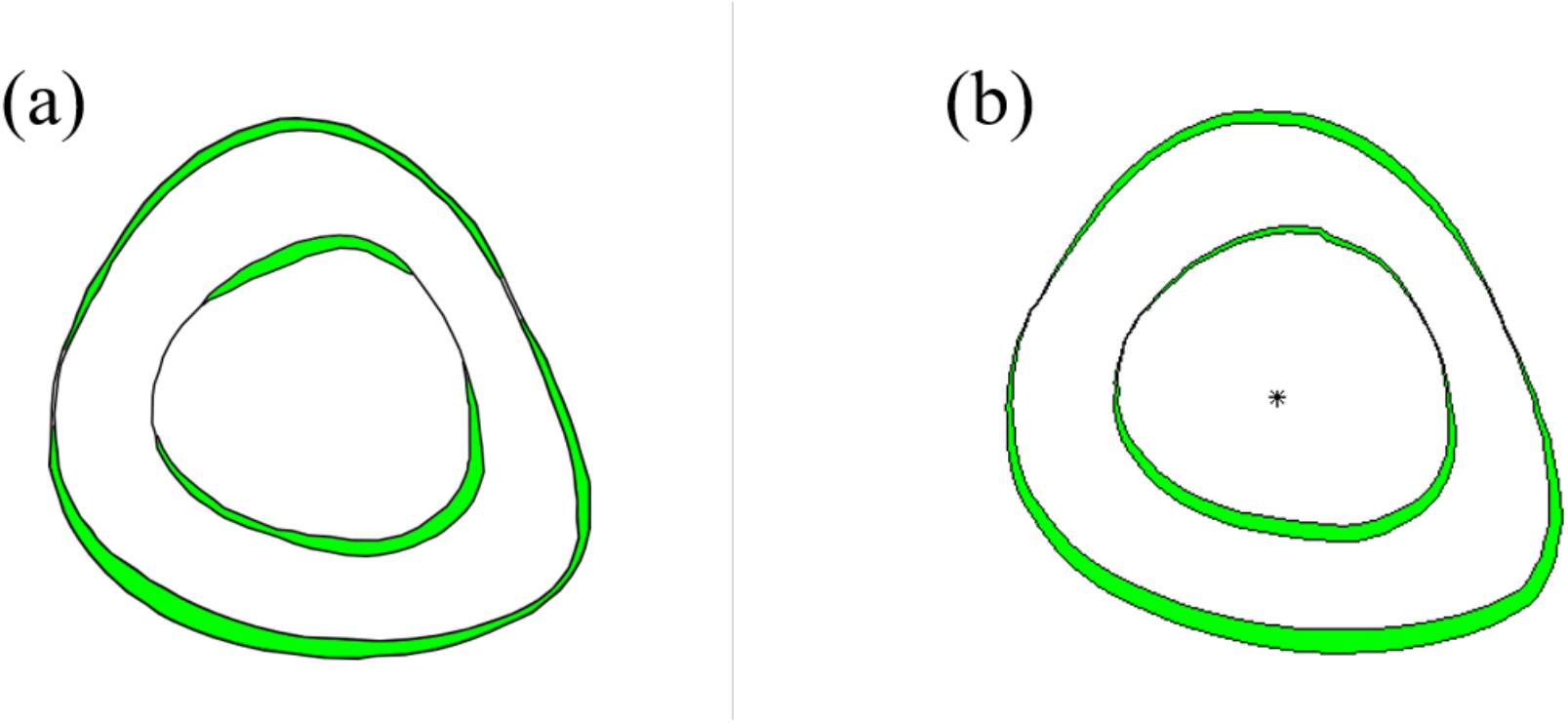
(a) In vivo new bone distribution at cortical envelops for high strain loading protocol (adapted from Berman et al. [22]) and corresponding computationally estimated new bone distribution using (b) a simplified model.

## 4 Sensitivity Analysis

In modeling studies, knowing how different factors affect the results is essential. In this study, we tuned the parameter, *A*and *ψ*_*ref*_ of a mathematical model while considering the value of *n* = 0.5 to predict the new bone distribution. To understand the physical significance of parameter *A*, and *ψ*_*ref*_, the simplest form of sensitivity analysis is used. The analysis is called local sensitivity analysis (LSA) or one-factor-at-a-time [51]. In this sensitivity analysis, we vary one model parameter at a time by a given amount and examine the impact on the output results.

Figure 10 (a) illustrates that increasing the remodeling rate (*A*) leads to a linear increase in the bone formation rate (*BFR*) while keeping the other parameter constant. Thus, we identify *A* as a parameter that amplifies the response of osteocytes. This lines up with another study done by Kumar et al. [48], who found that *A* influences how osteocytes react. As far as the dependence of *A* on loading parametres is concerned, Eq. (20) shows that *A* is a proportionality constant that does not depend on any loading parameters. On the other hand, Fig. 10 (b) demonstrates the impact of changing the parameter *ψ*_*ref*_ on the bone adaptation, while *A* remains specified. Interestingly, increasing the value of *ψ*_*ref*_ decreases the BFR linearly. *ψ*_*ref*_ may be identified as a threshold sensitivity of the osteocytes, which depends on the number of cycles (*N*) and days of loading (*d*). *ψ*_*ref*_ increases as the number of loading cycles increases (see Fig.10(c)) and saturates over time. This threshold sensitivity indicates osteocytes lose their sensitivity as the loading cycles increase. Other loading parameters, such as loading magnitude and frequency, do not change these tuning parameters as they directly influence the dissipation energy density per cycle.

**Fig. 10.**
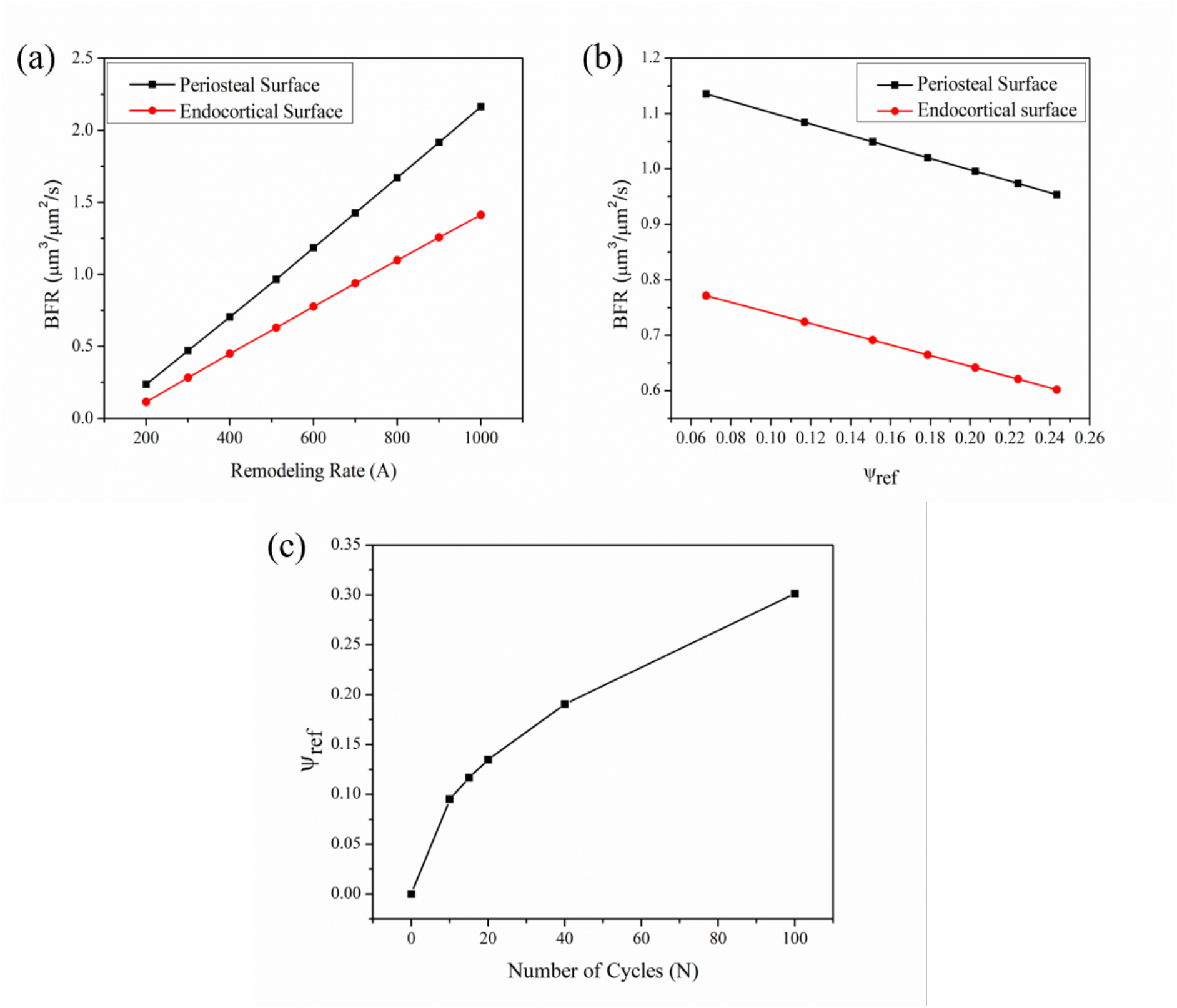
Plots showing the change in BFR to changing (a) remodeling rate (*A*) (b) threshold/reference (*ψ*_*ref*_).

## 5 Discussion

We utilize finite element analysis to determine the structural changes in the tibia mid-diaphyseal cross-section under the action of mechanical loading, and a novel mathematical formulation is derived and used to estimate the new bone formation at the cortical surfaces. The developed mathematical model exhibits that the combination of fluid flow with fluid pore pressure may be responsible for (re)modeling responses, i.e., MAR on both the periosteal and endocortical surfaces. A study by Tiwari et al. also confirms that two or more stimuli may be required to accurately predict osteogenesis at the cortical surfaces [49]. It is in contrast to the studies that consider a single stimulus, such as strain, velocity, etc., that finds success only in predicting bone formation at the periosteal surface [6, 40, 48]. Unlike previous mathematical models that measure the average bone formation rate at a cross-section [52, 53], our site-specific model provides a spatial distribution of new bone formed at both cortical surfaces, which gives our model an edge over the previous models.

In this work, we considered dissipation energy density due to fluid flow and pore pressure as a stimulus, as numerous studies suggest that they play a prominent role in bone adaptation [2, 6, 11, 19, 21]. The dissipation energy is the work done by the pore pressure or shear forces due to fluid flow experienced by the osteocytes’ membranes. The mathematical model results show that fluid flow alone as a stimulus predicts osteogenesis at the endocortical surface, whereas it underestimates the site-specific bone formation at the periosteal surface. The converse results were found when considering pore pressure alone as a stimulus, as it predicted bone formation at the periosteal surface and underestimated it at the endocortical surface. This was because of the drained and undrained conditions, respectively, of the endocortical and periosteal surface resulting in high fluid flow (or zero pressure) and negligible fluid flow (or high pressure), respectively, at the two surfaces (Fig. 7b, c) [21, 28]. This follows the literature where Carriero et al. show that fluid flow predicts the new bone distribution at the endocortical surface only. Along with it, Gardiner et al. and Scheiner et al. show that pore pressure also has the potential to act as a stimulus [7, 17, 21]. This suggests that fluid flow controls the new bone formation at the endocortical surface, whereas pore pressure controls the bone distribution at the periosteal surface.

This model tries to integrate most of the previously developed models. Cater et al. suggested a power relationship between bone adaptation and stress history, where the power exponent was an undetermined value and to be estimated by the quantitative analysis of in vivo studies [54]. Our model establishes a power law relationship between bone formation and stimulus with a power exponent 0.5. The total dissipation energy density (*φ*) is the function of other parameters, the number of cycles (*N*), and the number of days (*d*). The more cycles and days, the more will be the value of φ, which means more new bone formation; however, the osteogenic potential of the current cycle will be less than the previous cycle because of the power exponent *n*. This is in accordance with experimental and numerical literature that shows bone response saturates with the increase in the number of cycles [13, 40, 55, 56]. Similarly, the osteogenic potential of the current day’s loading will also be lower than the previous due to the exponent *n*. This also results in the saturation of MAR. However, the mechanism behind the saturation of MAR due to the number of cycles and days are different. The higher number of loading cycles results in the desensitization of osteocytes, resulting in the saturation of MAR. In contrast, the number of loading days increases the stiffness of bone, which leads to a reduction in stimulus (dissipation energy density). The previously developed poroelasticity-based models did not capture this aspect of bone adaptation as they have considered the bone adaptation to be linearly related to a stimulus (for example, Fluid velocity, dissipation energy density) [5, 6]. Note that similar to the stimulus (*φ*), the threshold/reference *ψ*_*ref*_ is also a function of number of cycles (*N*), and the number of days (*d*) of loading. In a way, it means that the threshold value of MAR also increases as osteocytes become “deaf” to repeated mechanical loading.

Bone formation and its spatial distribution are only tested on the midsection of the tibia. However, the model’s accuracy can be pushed to predict the new bone formation at different cross-sections along the tibial length. The model does not incorporate the erosion or resorption process during loading, which is generally detected near the neutral axis on the endocortical surface rather than the periosteal surface [57]. In addition, the developed model does not predict woven bone formation as the mechanism differs from the lamellar bone formation [58]. All the above cases, i.e., spatial distribution along the tibial length, erosion, and woven bone formation at one or both the periosteal and endocortical surfaces, have been taken as future work [59, 60]

The newly developed model made some simplifications concerning the structure and properties of the bone, which are as follows.

Structure – Bone has porosities at different lengths, such as lacune canalicular and vascular porosity [23], affecting bone formation rate. These vascular channels act as a local sink and show their shielding effect by reducing the fluid flow in its surrounding region [11, 61]. However, for simplification, we only considered the lacune canalicular porosities only. Moreover, we considered LCN porosity to be unrealistically regular, which, in reality, is spatially heterogeneous [62, 63]. The LCN architecture influences the fluid flow pattern and, consequently, its effect on osteocytes [11]. Incorporation of multiscale porosities may improve results.

Properties – The bone, for simplicity, has been considered a linear, isotropic, homogenous poroelastic material. However, the bone, in general, is anisotropic [64], which may affect the fluid flow in LCN [65]. Based on the existing limited literature, the permeability of the periosteal and endocortical surfaces, are assumed to be ideally permeable and impermeable, respectively. Our data (not included here) show that the permeability of periosteal and endocortical surfaces affects the fluid flow pattern. Therefore, more robust data on permeability at both cortical surfaces may be needed to accurately predict the fluid flow pattern.

Biological Factors – This newly developed tissue-level model focuses on the tissue-level response of bone adaptation and ignores the molecular and cellular elements, such as the role of integrins [16] and glycocalyx [66]. Moreover, biochemical molecules such as parathyroid hormone (PTH), insulin-like growth factors (IGFs), IP, etc., believed to be involved in transmitting signals from osteocytes to osteoprogenitor cells, have not been incorporated in this model to reduce the complexity [67].

## 6 Conclusion

To the authors’ best knowledge, a poroelasticity-based mathematical model presented here is the first of its kind to predict the site-specific lamellar bone distribution (MAR) simultaneously at both cortical surfaces (periosteal and endocortical). It shows that the site-specific mineral apposition rate is directly proportional to the square root of dissipation energy density due to both fluid flow and pore pressure minus its reference value. Analytical derivation of this relationship is also novel, and to the authors’ best knowledge, such derivation is being reported for the first time. The key finding of this model is that the bone formation rate at the periosteal and endocortical surface is primarily controlled by pore pressure and fluid flow, respectively. This novel model can be improved by integrating woven bone formation at higher loads and resorption at the endocortical surface. The model can be further improved by considering the effect of age.

## Acknowledgment

The author(s) would like to acknowledge IIT Ropar for providing facility and financial support to carry out the experimental test for the successful completion of the work.

## Nomenclature

LCN – Lacunae Canalicular Network BFR – Bone Formation Rate

*MAR*_*i*_ – Mineral apposition rate at the node of interest, µm^3^/µm^2^/day

*i* – Node of interest

*DE*_*v*_ – Dissipation energy density due to fluid flow per unit cycle, 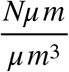

*DE*_*p*_ – Dissipation energy density due to pore pressure per unit cycle,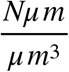

*DE*_*ref*_ – Reference/threshold value of dissipation energy density per unit cycle, 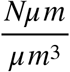

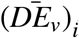– Modified dissipation energy density per unit cycle due to fluid flow at a given node after considering the zone of influence, 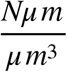

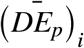 – Modified dissipation energy density per unit cycle due to pore pressure at a given node after considering the zone of influence, 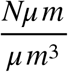

*n*_*p*_ – Porosity

***v*** ^*f l*^ – Fluid velocity at nodes, 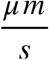

*P* – Pressure at the nodes, 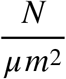

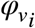– Total dissipation energy density due to fluid flow, 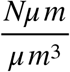

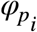– Total dissipation energy density due to pore pressure, 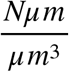

*ψ*_*ref*_ – Reference/threshold MAR in the absence of exogenous mechanical loading, µm^3^/µm^2^/day

*N* – Number of loading cycles

*d* – Number of loading days

